# Novel replication-competent reporter-expressing Rift Valley Fever Viruses for molecular studies

**DOI:** 10.1101/2024.08.06.606778

**Authors:** Aitor Nogales, Celia Alonso, Sandra Moreno, Gema Lorenzo, Belén Borrego, Luis Martinez-Sobrido, Alejandro Brun

**Affiliations:** Centro de Investigación en Sanidad Animal (CISA), Instituto Nacional de Investigación y Tecnología Agraria y Alimentaria, Consejo Superior de Investigaciones Científicas (INIA-CSIC), Valdeolmos, 28130 Madrid, Spain; Texas Biomedical Research Institute, San Antonio, TX, United States

**Keywords:** Rift valley fever virus (RVFV), segment S, reporter gene, NanoLuc luciferase, Venus fluorescent protein, neutralizing antibody, antiviral, mouse model

## Abstract

Rift Valley fever virus (RVFV) is a mosquito-borne zoonotic disease that causes severe disease in both domestic and wild ungulates and humans, making it a significant threat to livestock and public health. The RVFV genome consists of three single-stranded, negative-sense RNA segments differing in size: Small (S), Medium (M) and Large (L). Segment S encodes the virus nucleoprotein N and the virulence-associated factor non-structural (NSs) protein in opposite orientations, separated by an intergenic region (IGR). To overcome the current need of using secondary techniques to detect the presence of RVFV in infected cells, we used T7-driven polymerase plasmid-based reverse genetics to generate replication-competent recombinant (r)RVFV expressing Nanoluciferase (Nluc) or Venus fluorescent proteins. These reporter genes were used as valid surrogates to track the presence of RVFV in mammalian and insect cells. Notably, we explored the genome plasticity of RVFV and compared four different strategies by modifying the viral segment S in order to introduce the reporter gene foreign sequences. The reporter-expressing rRVFV were stable and able to replicate in cultured mammalian and insect cells, although to a lesser extent than the recombinant wild-type (WT) counterpart. Moreover, rRVFV expressing reporter genes were validated to identify neutralizing antibodies or compounds with antiviral activity. *In vivo*, all mice infected with the reporter-expressing rRVFV displayed an attenuated phenotype, although at different levels. These rRVFV expressing reporter genes provide a novel approach to better understand the biology and pathogenesis of RVFV, and represent an excellent biotechnological tool for developing new therapeutics against RVFV infections.

**IMPORTANCE:** Rift Valley fever virus (RVFV) is a mosquito-borne virus and zoonotic agent threat that can be deadly to domestic or wild ungulates, and humans. In this work, we used reverse genetics approaches to explore the genome plasticity of RVFV by generating a set of recombinant rRVFV that express fluorescent or luminescent proteins to track viral infection. All the generated reporter-expressing rRVFV were able to propagate in mammalian or insect cells, and in a mouse model of infection. Our studies may contribute to advances in research on RVFV and other bunyaviruses and pave the way for the development of novel vaccines and the identification of new antivirals for the prophylactic and therapeutic treatment, respectively, of RVFV infections.

## INTRODUCTION

Rift Valley fever (RVF) is a re-emerging zoonotic disease endemic to Africa, the Middle East, and the Indian Ocean with recurrent outbreaks in domestic or wild ruminants resulting in spillover into humans (1–3). Rift Valley fever virus (RVFV) is a phlebovirus belonging to the family *Phenuiviridae* that is transmitted by *Aedes* and *Culex* mosquito species, but may also be transmitted through direct contact with body fluids of infected animals (2). Human infections in endemic regions are usually asymptomatic or can cause mild illness. However, in some cases, infected persons develop more severe symptoms that range from ocular lesions to encephalitis and haemorrhagic fever or death (1). Moreover, RVF disease leads to high abortion rates in herds and neonatal mortality, making it a significant threat to livestock of affected regions (4). Due to its virulence, economic impact and lack of efficient vaccines or antivirals, RVFV is considered a potential biothreat and is listed as a select agent by both the U.S. Health and Human Services (HHS) and the U.S. Department of Agriculture (USDA). In addition, RVFV is included in the World Health Organization (WHO) Blueprint list of pathogens that are prioritised for research and development of countermeasures (5). Therefore, research to better understand RVFV pathogenesis and to develop effective prophylactic vaccines and therapeutic strategies is urgently needed for the control of this important human viral pathogen.

RVFV virions are icosahedral and contain a lipid envelope enclosing a nucleocapsid. The viral genome is formed by three single-stranded RNA segments with negative and ambisense polarity and different sizes: the large (L), medium (M), and small (S) (6, 7). The 3′ and 5′ ends of the coding regions contain regulatory untranslated regions (UTRs). The L segment encodes the RNA-dependent RNA polymerase (RdRp) (8, 9). The M segment encodes a polyprotein, which, upon protease processing, gives the two membrane glycoproteins (Gn and Gc) that form heterodimers in the viral envelope (10–13), and a non-structural protein (NSm) involved in modulation of apoptosis (14). The mRNA of the M segment may also translate an additional 78 kDa protein via the NSm and Gn open reading frames (ORFs) (15). The S segment encodes the virus nucleoprotein (N) and the virulence-associated factor non-structural (NSs) protein in opposite orientations, separated by an intergenic region (IGR) (16–18). Together with the viral RNA, the N and RdRp proteins constitute the viral replication complex. While the genomic L and M RNAs serve as templates for the synthesis of their cognate antigenomic RNAs and mRNAs, the segment S uses an ambisense coding strategy for the expression of N and NSs proteins in opposite orientation. The genomic S segment serves as a template for the synthesis of N mRNA and the antigenomic S segment serves as the template for the synthesis of NSs mRNA.

Reverse genetics methods have provided researchers with a powerful experimental approach for generating recombinant RVFV (rRVFV), which have allowed researchers to study important aspects of the biology and replication of RVFV, identify host factors, study markers of pathogenesis, and generate live-attenuated vaccines (19–28). The generation of reporter-expressing rRVFV has been described previously and engineered to contain four-segments: an authentic L segment, an S segment that encodes N and enhanced green fluorescent protein (eGFP), and two M-type segments that encode either NSmGn or Gc (29). This rRVFV-4s eGFP was able to reach high titters in mammalian BSR cells, but not in insect C6/36 cells (29). Moreover, the virus was highly attenuated in mice (29). In addition, the generation of rRVFV lacking NSs (ΔNSs) expressing either eGFP or firefly luciferase (FLuc) in place of NSs have also been described (22). The growth curves of recombinant wild-type (WT) and ΔNSs-eGFP or ΔNSs-FLuc were similar in Vero cells, but the growth of the reporter viruses in mice was highly attenuated compared to their WT counterpart (22). Moreover, while the eGFP sequence was well tolerated in the viral genome, FLuc sequence was unstable when expressed by the rRVFV (22).

In this work we used a new experimental strategy to generate replication-competent, reporter-expressing rRVFV in which heterologous genes are inserted into the viral genome without the need for segment duplication or gene deletion. Notably, we used four different approaches that also provided information related to the genome plasticity of RVFV. While numerous reporter genes with different features have been described, fluorescent and bioluminescent proteins are becoming the preferred choice for researchers due to their high sensitivity and the continuous improvement of the technologies associated with their detection (30–34). Therefore, we have generated a set of rRVFV expressing fluorescent (Venus) or nanoluciferase (Nluc) reporter genes. All viruses were characterized *in vitro* and *in vivo* and proven to be a valid method for studying viral replication and propagation or for the identification of antivirals and neutralizing antibodies (NAbs).

## RESULTS

### Generation of replication-competent rRVFV expressing reporter genes

To generate replication-competent, reporter-expressing rRVFV (strain 56/74), the sequence of Nluc or Venus and the porcine teschovirus 1 (PTV-1) 2A autoproteolytic cleavage site were cloned upstream or downstream of the ORF of the N or NSs proteins in the segment S (**Figure 1**). We chose Nluc due to its small size (∼20 kDa), stability, and brightness (30, 35). The fluorescent protein Venus (∼26,8 kDa) was also selected because of the intensity of the signal and previous experience with other RNA viruses, where the insertion of Venus and Nluc in the genome was well tolerated (36–39). Importantly, the UTRs and the IGR of the S segment were not modified. We then used T7-pol plasmid-based reverse genetics approaches to generate a set of rRVFV expressing Nluc or Venus at different genome positions. Notably, 4 Nluc-expressing rRVFV, covering the 4 strategies used, were recovered, namely Nluc2ANSs, NSs2ANluc, N2ANluc, and Nluc2AN. However, only 3 Venus-expressing rRVFV were rescued: Venus2ANSs, NSs2AVenus, and Venus2AN (**Figure 1**). We were not able to successful rescue the N2AVenus even after multiple attempts. Moreover, a rRVFV WT (rWT) was also generated for comparison.

**Figure 1.**
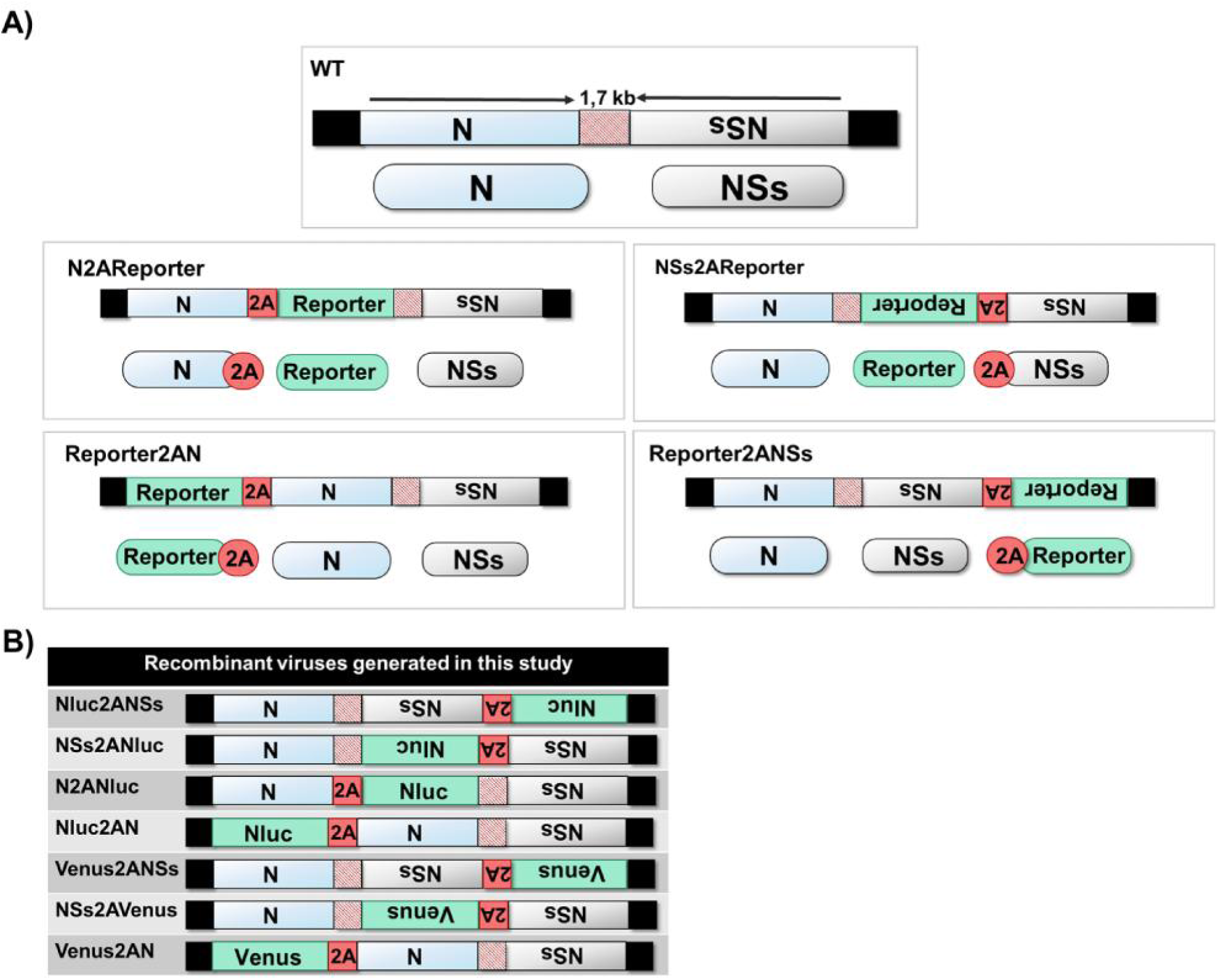
Schematic representation of the RVFV segment S. RVFV segment S and its viral products are indicated by blue (N) or gray (NSs) boxes. The black and striped red boxes denote untranslated regions and intergenic regions, respectively. The sequences of the reporter and 2A are indicated in green and red boxes, respectively.

### Growth properties and plaque phenotype of reporter-expressing rRVFV in mammalian cells

We next evaluated the fitness of Nluc-expressing (**Figures 2A and B**) or Venus-expressing (**Figures 2C and D**) rRVFVs in mammalian cells by assessing their growth properties and compare them to those of rWT. To this end, confluent monolayers of Vero E6 cells were infected with the different viruses at the same multiplicity of infection (MOI; 0.01), and the presence of virus in culture supernatants was quantified at different times post-infection (p.i).

**Figure 2.**
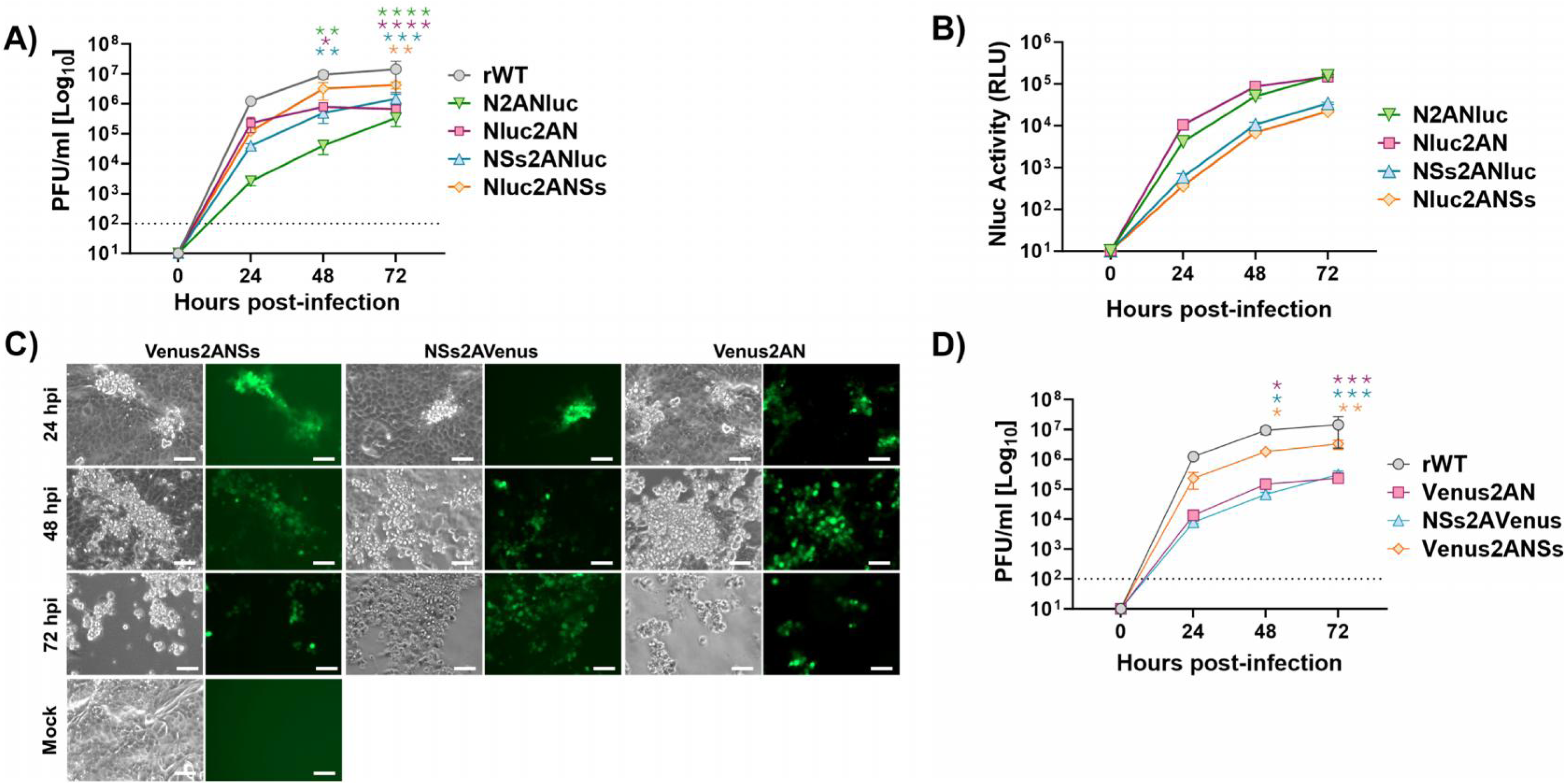
Multicycle growth kinetics. Vero E6 cells (12-well plates, triplicates) were infected (MOI of 0.01) with rRVFV WT (rWT) or with recombinant viruses expressing Nluc (**A and B**) or Venus (**C and D**). At the indicated h.p.i. (24, 48 and 72), viral titers in culture supernatants were evaluated by plaque assay (PFU/ml), and the same rWT representation was used in A and D (A and D). Comparison of the replication curves of viruses expressing Nluc or Venus versus rWT at different post-infection times revealed statistically significant differences (Dunnett’s method). * P < 0.05; ** P < 0.0021; *** P < 0.0002; **** P < 0.0001. The data represent the means ± SDs of triplicate samples. At the same time, post-infection Nluc activity was evaluated (RLU: Relative Light Units) (**B**), and Venus expression was analyzed using a fluorescence microscope (**C**). Representative images are shown. Scale bars, 100 μm. Dotted line indicates the limit of detection (100 FFU/ml) of the assay.

Replication of Nluc-expressing rRVFVs was delayed and did not reach titters similar to those of the rWT (**Figure 2A**). Moreover, higher replication levels were observed when Nluc was inserted next to the UTRs (Nluc2AN and Nluc2ANSs), instead of next to the IGR (N2ANluc and NSs2ANluc) and when the reporter gene was fused to the NSs ORF (Nluc2ANSs and NSs2ANluc) instead of the N ORF (Nluc2AN and N2ANluc) (**Figure 2A**). In addition, Nluc activity in culture supernatants was evaluated at the same time points (**Figure 2B**). Nluc activity increased in a time-dependent manner, peaking at 72 h p.i. (the last time point evaluated in our study), most likely because the cytopathic effect (CPE) caused by the viral infection resulted in the release and accumulation of Nluc in the cell culture supernatants. Interestingly, viruses expressing Nluc in the same ORF that the viral N gene (Nluc2AN and N2ANluc) displayed higher levels of Nluc activity (**Figure 2B**). This is most likely because of the higher levels of N expression during viral infection as compared to NSs (40).

Similarly, the viral growth of Venus-expressing rRVFVs was delayed as compared with that of rWT, and the replication of Venus2AN was more affected than the replication of Venus2ANSs (**Figures 2C and D**). Venus expression was detected as early as 24 h p.i. and steadily increased, in a time-dependent manner, until 72 h p.i., when visualization was reduced due to the virus-induced CPE. Importantly, all reporter-expressing rRVFVs were still able to reach high viral titters in culture supernatants (∼10^5/6^ PFU/ml) but in all cases were reduced as compared to rWT (∼10^7^ PFU/ml) (**Figure 2**). Our data suggest that the insertion of Nluc in the viral genome is better tolerated than the insertion of Venus. In addition, our results also provide some clues regarding the regions of RVFV segment S that can be more suitable for the expression of foreign sequences. Therefore, results indicate that reporter-expressing rRVFVs may be used to track and quantify the dynamics of viral infection in vitro by using either fluorescent or luciferase reporter genes.

We also evaluated the plaque phenotype of the rRVFVs in Vero E6 and compared it to the rWT virus (**Figure 3**). For that, Vero E6 cells were infected and at 72 h p.i., immunostaining using specific antibodies against Nluc or GFP (for Venus detection), and crystal violet staining. As expected, we detected Nluc or Venus-positive viral plaques only in cells infected with Nluc- or Venus-expressing rRVFVs, respectively. In addition, all the Nluc- or Venus-positive plaques colocalized with the plaques detected by crystal violet staining, confirming that all the rRVFV-infected plaques expressed the Nluc or Venus reporter gene. Moreover, plaque sizes were in agreement with viral replication kinetics (**Figure 2**), since the sizes of the viral foci produced by reporter-expressing rRVFVs were smaller than those of the produced by the rWT virus (**Figure 3**), most likely due to the effect of the reporter gene on viral fitness. Overall, these data and our previous studies with other reporter-expressing replication-competent viruses (36, 38, 41, 42) demonstrate that Nluc and Venus can be successfully expressing from the viral genome using the PTV-1 2A site, and reporter gene expression can be used as valid surrogates to monitor viral infection without the need for secondary approaches to detect the presence of the virus in infected cells, or the presence of virus in culture supernatants.

**Figure 3.**
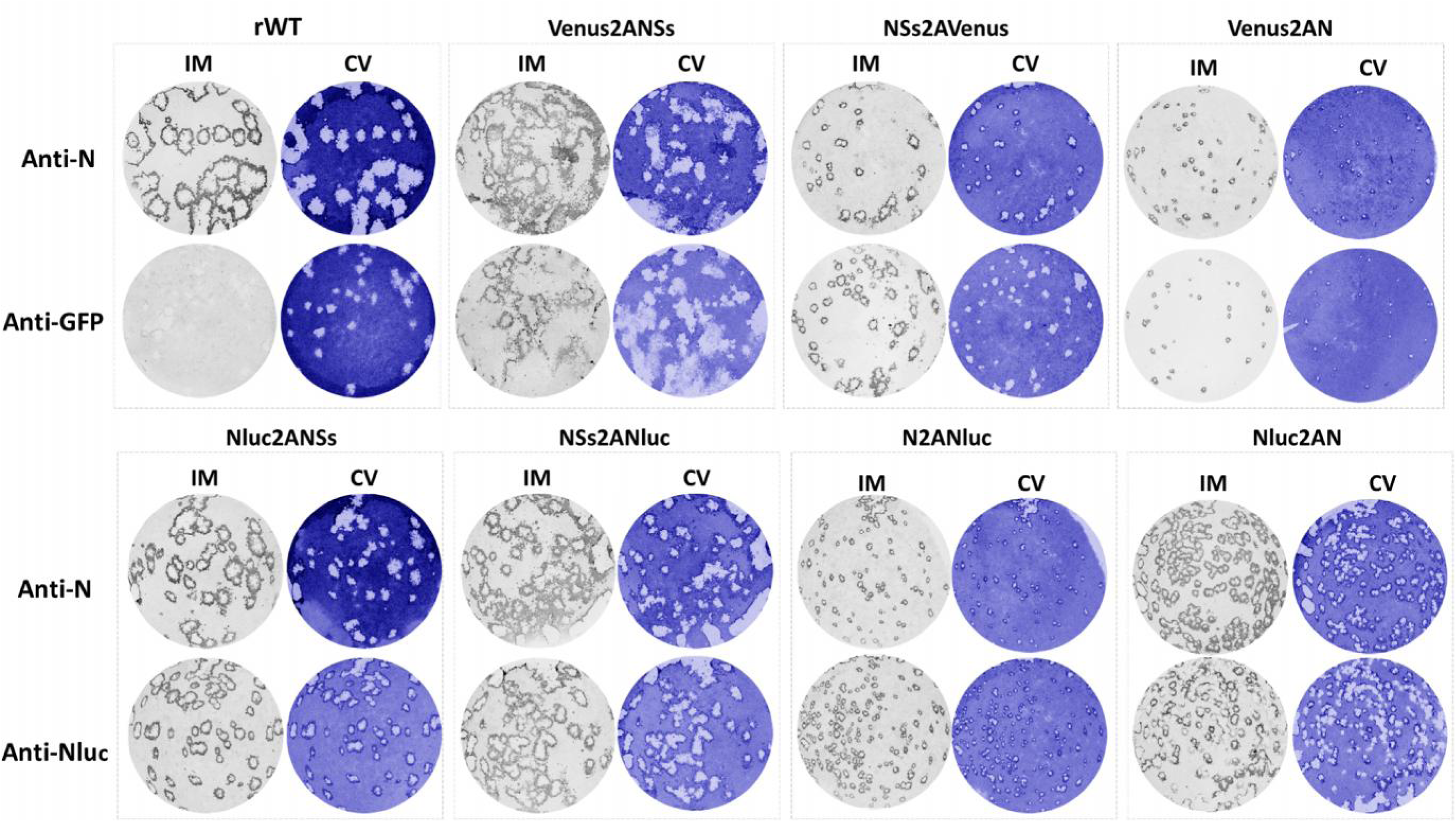
Plaque assays. Vero E6 cells (10^6^ cells/well, 6-well plate format) were infected with rRVFV WT (rWT) or with recombinant viruses expressing Venus (top) or Nluc (bottom) and incubated at 37°C for 3 days. Plaques were evaluated by immunostaining with the mAb anti-N (2B1), the goat pAb anti-GFP (top) or the mAb anti-Nluc (bottom). Immunostaining of rWT was performed with the mAb anti-N (2B1) or a combination of the goat pAb anti-GFP and the mAb anti-Nluc.

### Analysis of protein expression in cells infected with reporter-expressing rRVFVs

To assess the expression of Nluc and Venus, confluent monolayers of Vero E6 cells were mock-infected or infected with rWT and Nluc-expressing rRVFVs (**Figure 4A**) or Venus-expressing rRVFVs (**Figure 4B**). Then, the cells were analyzed by indirect immunofluorescence microscopy and confocal microscopy, using antibodies against RVFV (detecting Gn/Gc and N viral proteins), Nluc (**Figure 4A**), or GFP (for Venus detection) (**Figure 4B**). As expected, only rRVFV-infected cells were stained with the anti-RVFV polyclonal antibody (pAb). Accordingly, the expression of Nluc and Venus was detected only in cells infected with Nluc- (**Figure 4A**) or Venus-expressing (**Figure 4B**) rRVFVs, respectively. Interestingly, Nluc was detected in the cytosol resembling the characteristic intranuclear NSs filaments in both Nluc2ANSs and NSs2ANluc viruses, suggesting that the cleavage efficiency of the 2A peptide sequence selected was not 100%, as previously described (43, 44) and shown in figure 5. In contrast, the Nluc signal colocalizes along the RVFV signal in the cytoplasmic compartment in cells infected with either N2ANluc or Nluc2AN viruses, most likely in these cells some fraction of Nluc is also colocalizing with the viral N protein. The presence of intranuclear NSs accumulation was barely detectable in both NSs2AVenus and Venus2ANSs viruses in which both RVFV and Venus signals colocalize in the cytoplasmic compartment as of Venus2AN virus.

**Figure 4.**
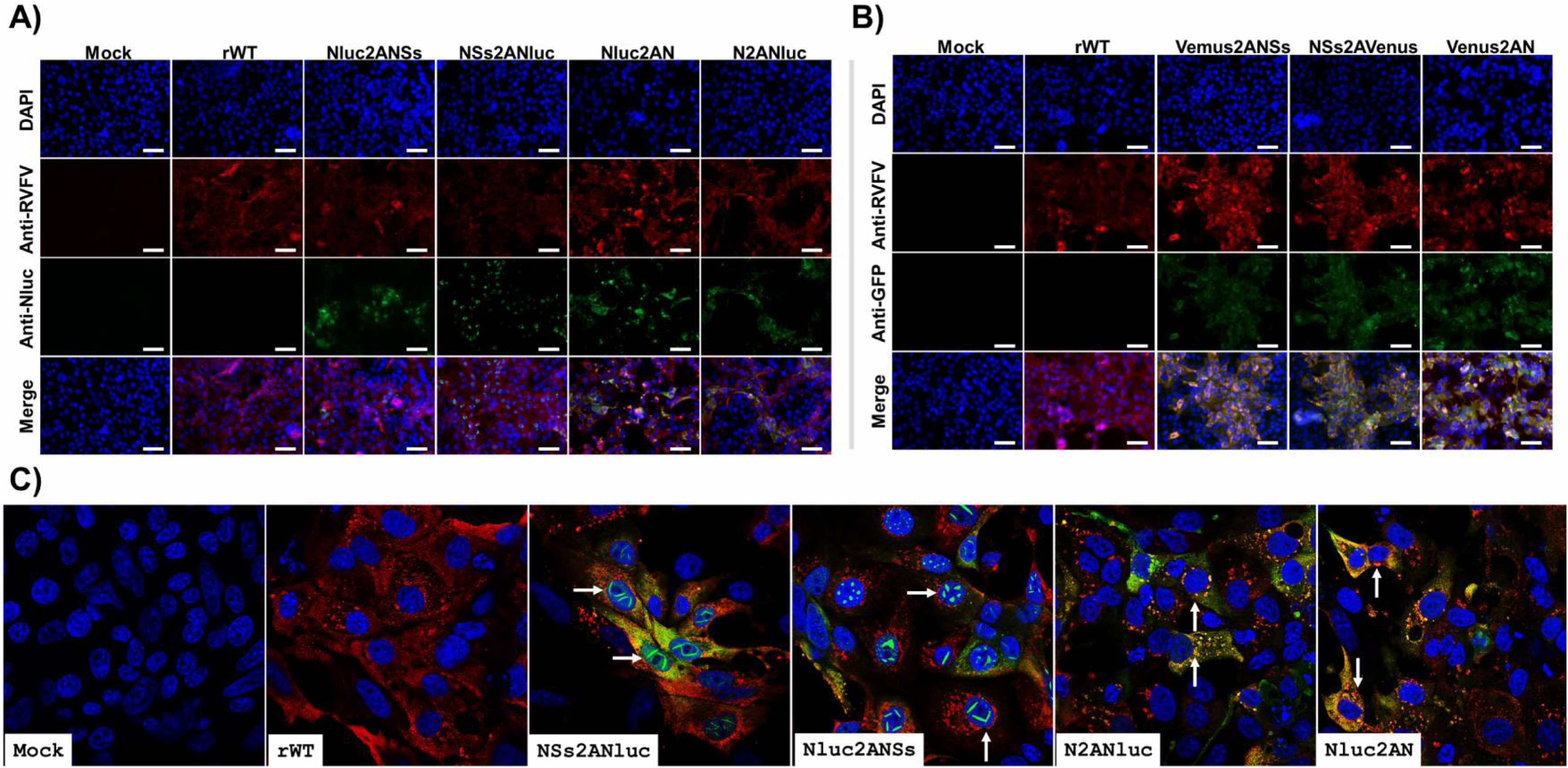
Analysis of protein expression by IFA. Vero E6 cells were infected with the indicated virus at an MOI of 0.1 or mock infected. At 24 h p.i., the cells were fixed and permeabilized. Protein expression was examined by IFA using a rabbit pAb against RVFV or mAb for Nluc (**A**) or a goat pAb against GFP (**B**). Scale bars, 100 μm. (**C**) **Confocal microscopy of Nluc-expressing rRVFV.** Protein expression was examined by confocal microscopy using the indicated antibodies. Arrows shows the presence characteristic intranuclear NSs filaments (Nluc2ANSs and NSs2ANluc) and colocalization of Nluc with viral proteins (Nluc2AN and N2ANluc).

**Figure 5.**
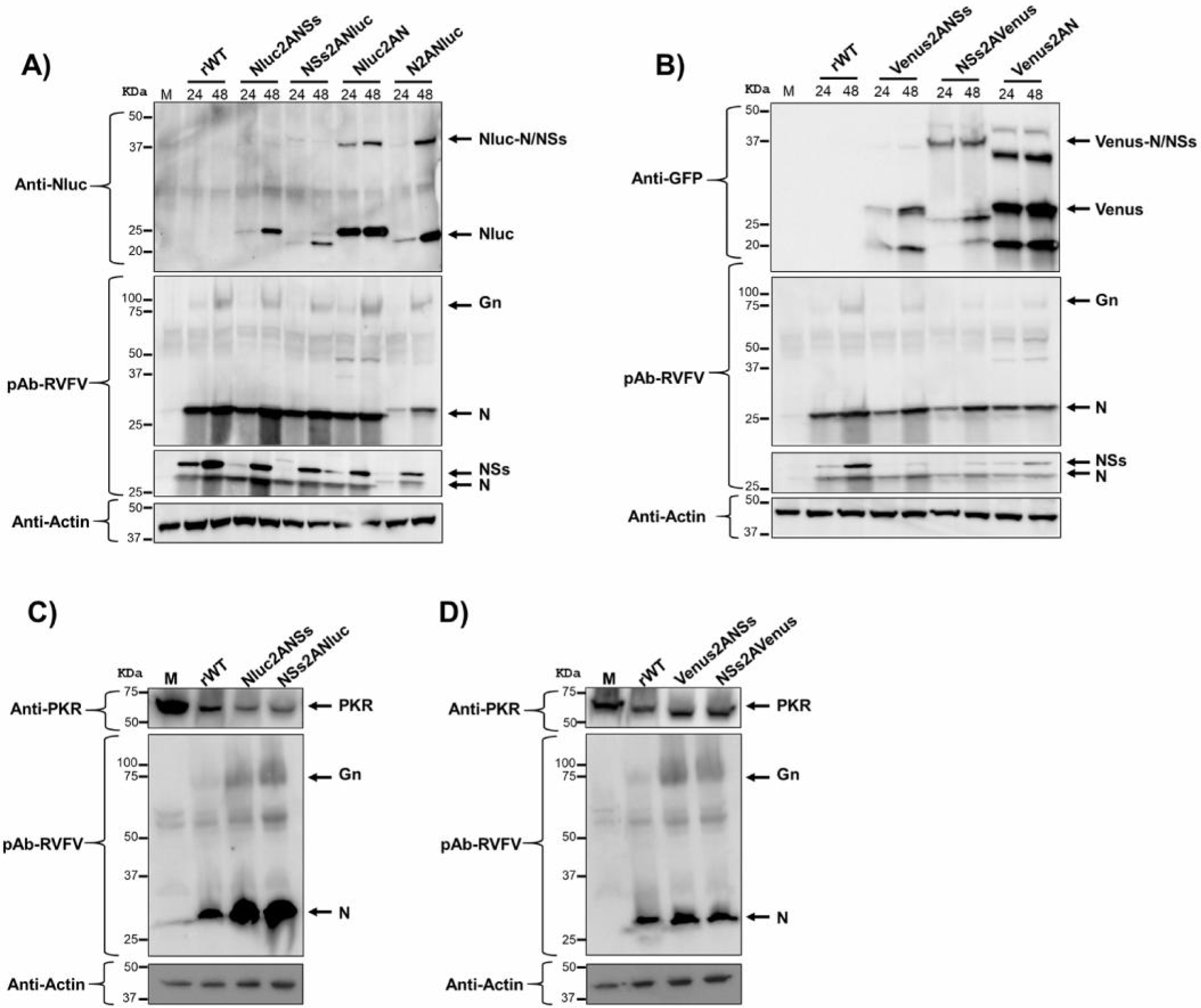
Analysis of protein expression by WB. Vero E6 cells were infected with Nluc- (**A**) or Venus-expressing (**B**) rRVFV at an MOI of 0.01 or mock infected. At 24 or 48 h p.i, cell extracts were prepared in RIPA buffer. Protein expression was examined by Western blotting using specific antibodies for viral proteins: pAb against RVFV for N and Gn, a pool of mAbs against NSs (after stripping of pAb). In addition, the expression of the reporter genes Nluc (**A**) and Venus (**B**) was evaluated. Actin was used as a loading control. The numbers on the left indicate the molecular size of the protein markers (in kilodaltons). (**C** and **D**) **Degradation of PKR in rRVFV infected cells.** HEK293T cells were infected at a MOI of 0.1 with the indicated viruses. Cells were harvested at 24 hpi and analyzed by Western blot using a pAb anti-PKR, a pAb against RVFV for N and Gn/Gc, and an anti-actin antibody as primary antibodies.

Next, protein expression in Vero E6 cells was evaluated by Western blot at 24 and 48 h p.i. (**Figure 5**). Total cell extracts from either mock-, rWT-, or reporter-expressing rRVFV-infected Vero E6 cells were tested using antibodies specific for RVFV, N or NSs, Nluc (**Figure 5A**) or Venus (**Figure 5B**) reporters, and for the loading control, actin. Western blot analysis revealed specific bands corresponding to the expected molecular sizes of the viral Gn (54 kDa) and Gc (56 kDa), N and NSs proteins. The N protein expressed by N2ANluc virus displays a slightly higher molecular mass as a consequence of the proteolytic mechanism that leaves the linker and part of the 2A peptide sequence at the C-term of the fusion partner. Interestingly, this shift is also observed for NSs (NSs2ANluc) although only at 24 h p.i. In addition, we observed specific bands for Nluc or the fusion protein of Nluc with NS or N (**Figure 5A**) and for Venus or the fusion protein Venus with NS or N (**Figure 5A**), indicating that the proteolytic efficiency of the 2A peptide sequence used was not 100%, as previously described (43, 44). Moreover, protein expression was greater at 48 h p.i. In the case of cell extracts infected with Venus-expressing rRVFVs, we also observed bands of lower molecular size than expected, which could indicate Venus degradation.

A pathway by which RVFV NSs blocks the host antiviral responses of infected cells is the degradation of cellular proteins such as protein kinase R (PKR) (45–47). In order to determine whether our gene modifications affected the ability of modified NSs to degrade PKR and therefore be responsible of viral attenuation, HEK293T cells were infected with rRVFV expressing the reporter gene in the NSs locus (Nluc2ANSs, NSs2ANluc, Venus2ANSs, and NSs2AVenus), and whole cell extracts were analyzed by Western blot (**Figure 5C**). In all infected cells, the expression of PKR was reduced, suggesting that the viral NSs is functional in all cases or at least its ability to degrade PKR was not impaired.

### Intracellular *vs* extracellular activity of Nluc and analysis of intracellular Venus expression

To determine whether differences in Nluc activity in cell culture supernatants were due to differences in the levels of Nluc secretion *versus* its accumulation in infected cells, confluent monolayers of Vero E6 cells were infected (MOI of 0.01) with Nluc-expressing viruses, and the presence of virus in culture supernatants was evaluated at 48 h p.i. (**Figure 6A**). In addition, Nluc activity in the supernatant or in cell extracts was quantified (**Figure 6B**). There was a correlation between Nluc activity in the supernatants and that in the cell extracts, suggesting that differences in Nluc activity were not a consequence of a greater secretion into the supernatant or cell accumulation of Nluc; indeed, these differences were most likely due to differences in the Nluc position in the viral genome. However, the observed intranuclear detection of Nluc aggregates (**Figure 4**), resembling NSs filamentous structures, might correlate with lower availability of soluble Nluc in cells infected with the NSs2ANluc expressing virus.

**Figure 6.**
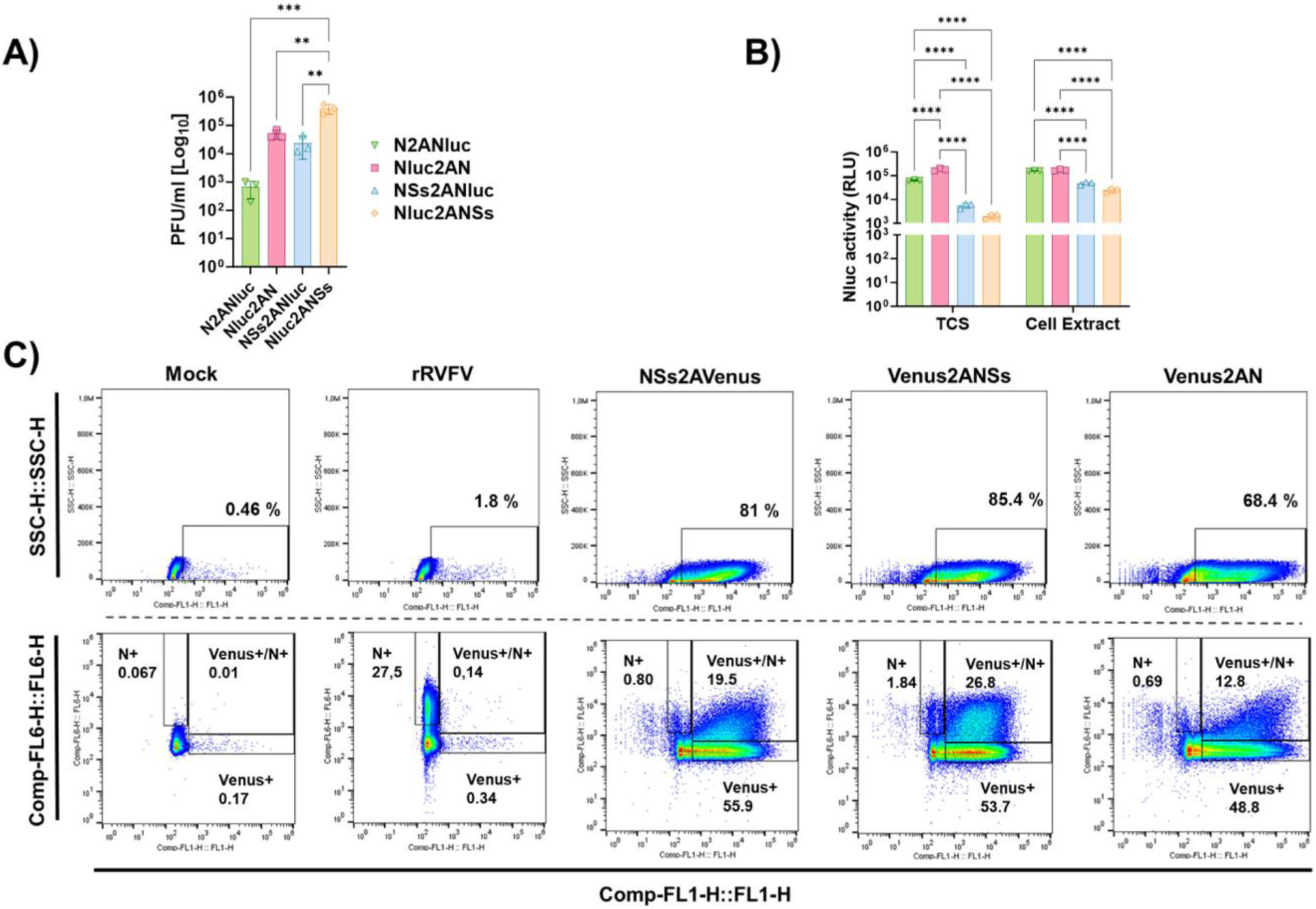
Nluc activity in tissue culture supernatant *versus* cell extracts and Venus expression by Flow cytometry analysis. Vero E6 cells (24-well plates, triplicates) were infected (MOI of 0.01) with recombinant viruses expressing Nluc. At 48 h p.i., viral titers in culture supernatants were evaluated by plaque assay (PFU/ml). The differences between the groups were calculated using one-way ANOVA with the Tukey method. * P < 0.05; *** P < 0.0002; **** P < 0.0001 (**A**). Nluc activity in tissue culture supernatants (TCSs) and cell extracts was measured. RLU: Relative Light Units. The differences between the groups were calculated using two-way ANOVA with the Tukey method. * P < 0.05; *** P < 0.0002; **** P < 0.0001. (**B**). The data represent the means ± SDs of triplicate samples. (**C**) Flow cytometry analysis. Vero E6 cells were infected (MOI of 0.01) with recombinant viruses expressing Venus. At 24 h p.i., the cells were harvested from the culture and stained with a mAb against N conjugated with Alexa 700. The cells were analyzed by flow cytometry by gating on green (Venus) and red fluorescence (mAb anti-N 2B1) and the percentage of N+, Venus+ or Venus+/N+ is indicated.

Next, Vero E6 cells infected with Venus-expressing rRVFVs were evaluated by flow cytometry to quantify the percentage of cells expressing Venus, N protein, or Venus and N protein at 24 h. p.i, using the anti N monoclonal antibody (mAb) 2B1conjugated to Alexa 674 (**Figure 6C**). Notably, the percentage of Venus+ cells was greater than the percentage of N+ cells. Moreover, 100% of the Venus+ cells were also N+. In contrast, and as expected, the rWT infection control was detected only after N staining. Our data indicate that the expression of Venus during viral infection could be a more sensitive method than antigen staining to detect the presence of the virus in infected cells, even more than detecting one of the highly expressed proteins during viral infection (e.g. N), at least, at early times post infection.

### *In vitro* stability of the reporter-expressing rRVFV

The genetic stability of reporter-expressing viruses is important for their use since can limit some of their applications (36, 38). Therefore, we evaluated the *in vitro* stability of all the reporter-expressing rRVFVs generated (**Figure 7**). To that end, Nluc- or Venus-expressing rRVFVs were passaged in duplicate three consecutive times in Vero E6 cells, and the percentage of reporter-expressing viruses was determined by plaque assay. In the case of Venus-expressing viruses, plaques were observed directly under a fluorescence microscope. To test the expression of Nluc, plaques were amplified in 96-well plates, and then, Nluc activity in the cell culture supernatant was evaluated at 48 h.p.i. rRVFVs that express Nluc or Venus were 100% stable at passage two (virus working stock) and passage 3 (first amplification passage) in all cases. Interestingly, the rRVFVs expressing the reporter genes close to the IGR (NSs2ANluc, N2ANluc, and NSs2AVenus) were more stable than the viruses encoding Nluc or Venus next to the UTRs (Nluc2ANSs, Nluc2AN, and Venus2ANSs), where a gradual loss of Nluc or Venus expression was observed from passages 5 or 4, respectively (**Figure 7**). Venus2AN was an exception since it expressed stable fluorescent levels up to 5 passages.

**Figure 7.**
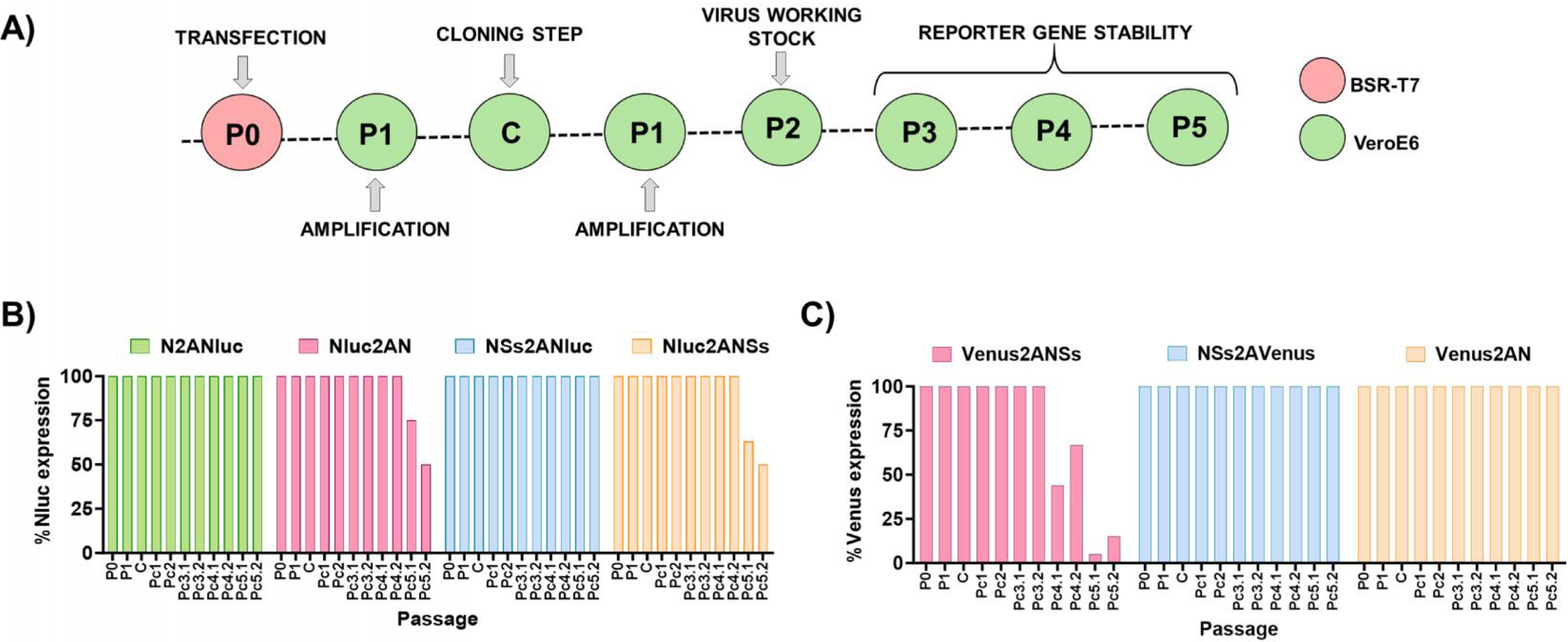
Genetic stability of reporter-expressing RVFVs in vitro. (**A**) After the generation of virus working stocks (P2), the recombinant viruses were passaged up to 3 times in Vero E6 cells (P3 to P5). Stability is expressed as the percentage of plaques that expressed Nluc (**B**) or Venus (**C**) reporter genes in each passage.

### Use of rRVFV expressing Nluc for the identification of antivirals

Identification of compounds that inhibit RVFV normally requires the use of secondary approaches to evaluate viral infection (48–51). The use of reporter-expressing viruses compatible with high-throughput screening for the identification of antivirals may be an interesting alternative to traditional assays. To demonstrate that Nluc-expressing rRVFVs are a valid surrogate for identifying antivirals, we examined the ability of ribavirin to inhibit RVFV infection using the Nluc-expressing rRVFVs NSs2ANluc, Nluc2ANSs, and Nluc2AN (**Figures 8A and B**). We did not include the virus N2ANluc in the assay, because this rRVFV was the most affected in viral replication (**Figure 2**). At high concentrations of ribavirin (starting concentration of 1000 μM), all Nluc-expressing rRVFV were inhibited in a dose-dependent manner, as determined by Nluc activity in cell culture supernatants (**Figure 8A**). Importantly, the IC_50_ values were similar for all the viruses tested (**Figure 8B**) and consistent with those described in the literature (48–52).

**Figure 8.**
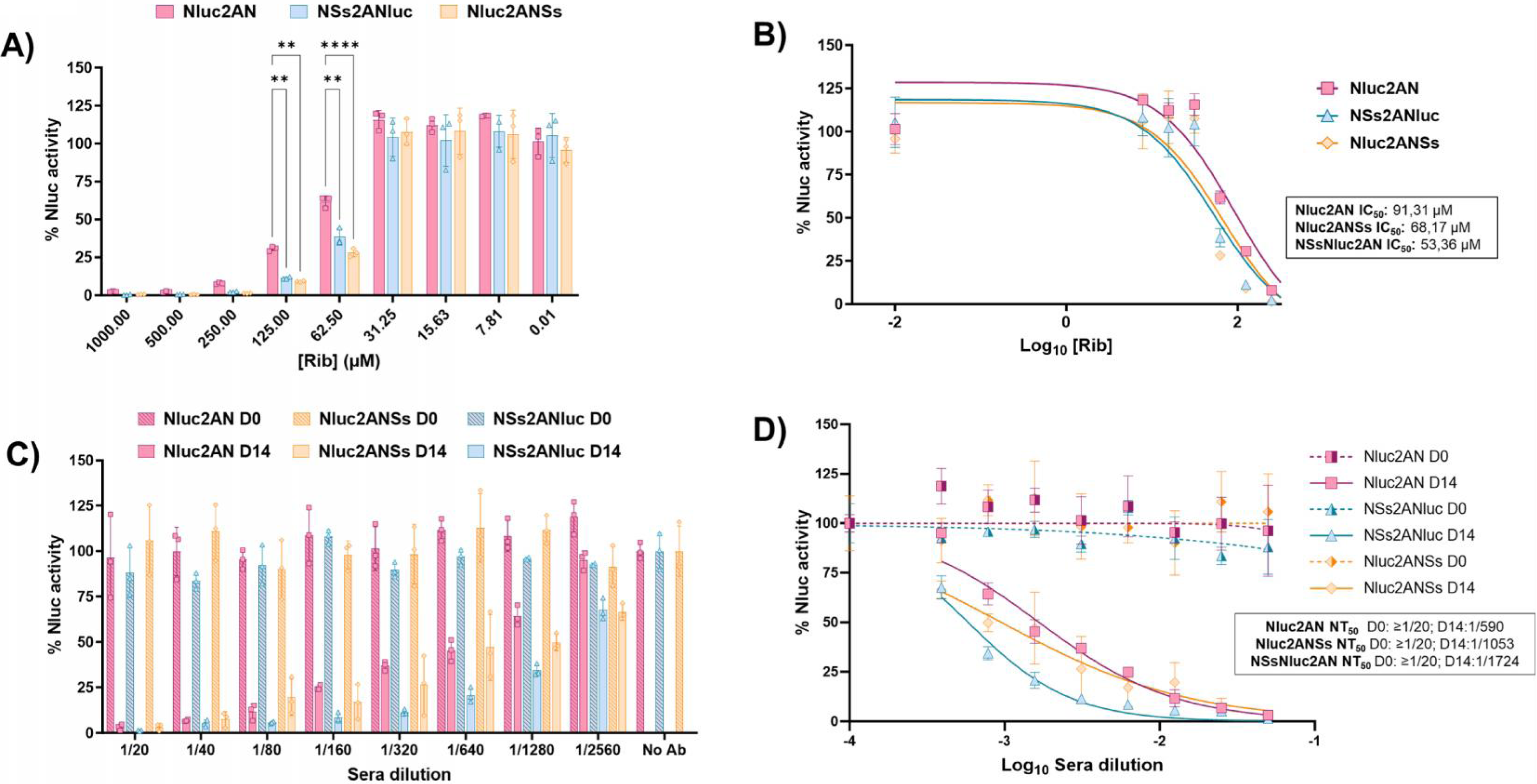
A luciferase-based microneutralization assay for assessing RVFV antivirals and neutralizing antibodies. (A and B) Antiviral. Vero E6 cells (96-well plates, 2 x 10^4^ cells/well, triplicates) were infected with 200 PFU of Nluc-expressing rRVFV and incubated with 2-fold serial dilutions (starting concentration, 1000 µM) of ribavirin. Inhibition of viral replication was evaluated by quantifying Nluc reporter expression at 48 h p.i. The differences between the groups were calculated using two-way ANOVA with the Tukey method. * P < 0.05; *** P < 0.0002; **** P < 0.0001. (**A**). The percent inhibition (IC_50_) was calculated using sigmoidal dose-response curves (**B**). As an internal control, infected cells were not treated with antivirals. The percentage of viral inhibition was normalized to that with infection in the absence of antivirals. The data are presented as the means ± SDs of the results determined in triplicate. (**C and D**) **Neutralizing antibodies.** Two hundred PFU of Nluc-expressing rRVFV were pre-incubated with 2-fold serial dilutions (starting dilution, 1:20) of sheep sera collected before or after vaccination for 1 h. Subsequently, Vero E6 cells (96-well plates, 2 x 10^4^ cells/well, triplicates) were infected with the sera-virus mixture. (**A**) Virus neutralization was determined by quantitating Nluc reporter expression at 48 h p.i. (**B**) The percent neutralization (NT_50_) was calculated using sigmoidal dose-response curves. Infected cells in the absence of serum and mock-infected cells were used as internal controls. The data are presented as the means ± SDs of the results determined in triplicate.

### Use of rRVFV expressing Nluc for the identification of NAbs

NAbs are a key immunological outcome for the induction of protective immunity after vaccination to prevent RVFV infections. Similar to the previously described antiviral test, viral neutralization assays typically involve the use of secondary methods to measure viral infection. Because the presence of Nluc in the culture supernatants is a valid surrogate of viral replication, an Nluc-based microneutralization assay was developed using the Nluc-expressing rRVFVs. Confluent monolayers of Vero E6 cells were infected with Nluc-expressing rRVFV, which had previously been incubated with sheep sera obtained before or at 14 days post-vaccination with an experimental RVFV vaccine (20) (**Figures 8C and D**). Then, at 48 h p.i. Nluc activity in the culture supernatants was quantified (**Figure 8C**). As expected, all viruses were neutralized, in a dose-dependent matter, by sera collected at 14 days post-vaccination. Using sigmoidal dose-response curves, we determined the 50% neutralization titter (NT_50_) obtained by the quantification of Nluc (**Figure 8D**). Importantly, the NT_50_ was similar for all the tested viruses. These data indicate that our Nluc-based-microneutralization assays may be a suitable method for the rapid identification and quantification of antivirals and NAbs.

### Growth properties of rRVFV in C6/36 mosquito cells

RVFV is an arbovirus, with mosquitoes playing an important role in the natural transmission cycle. Therefore, the growth properties of the reporter-expressing rRVFVs were also investigated in the *Aedes albopictus* clone mosquito larvae derived cell line (**Figure 9**). For that, C6/36 cells were infected (MOI; 0.01) with the different viruses, and the presence of virus in culture supernatants was quantified at 3, 5 and 7 days p.i. As expected, we observed differences in the levels of viral replication among the rRVFVs. Nluc-expressing rRVFVs replicated to a lesser extent than the rWT. In addition, as previously observed in Vero E6 cells (**Figure 2**), higher replication levels were observed when Nluc was cloned in the ORF of the NSs gene (Nluc2ANSs vs Nluc2AN or NSs2ANluc vs N2ANluc) (**Figure 9A**). Furthermore, the presence of Nluc in cell culture supernatants was measured at the same time points. Nluc activity increased in a time-dependent manner, peaking at 7 days p.i. (the last time point evaluated). Similar to Vero E6 cells, viruses expressing Nluc from the ORF of the viral N gene displayed also high levels of Nluc activity, most likely because of the higher levels of N expression during viral infection than NSs (40) (**Figure 9B**). In contrast to what was detected in Vero E6, the NSs2ANluc virus also had high Nluc activity level. The viral replication of Venus-expressing rRVFVs was also delayed as compared with that of the rWT (**Figure 9C and D**). Venus expression was detected as early as 3 days p.i. (first time point evaluated) and steadily increased in a time-dependent manner, until 7 days p.i. (last time point evaluated). Therefore, the data indicate that Nluc- and Venus-expressing rRVFVs may also be used to track the dynamics of viral infection in mosquito cells.

**Figure 9.**
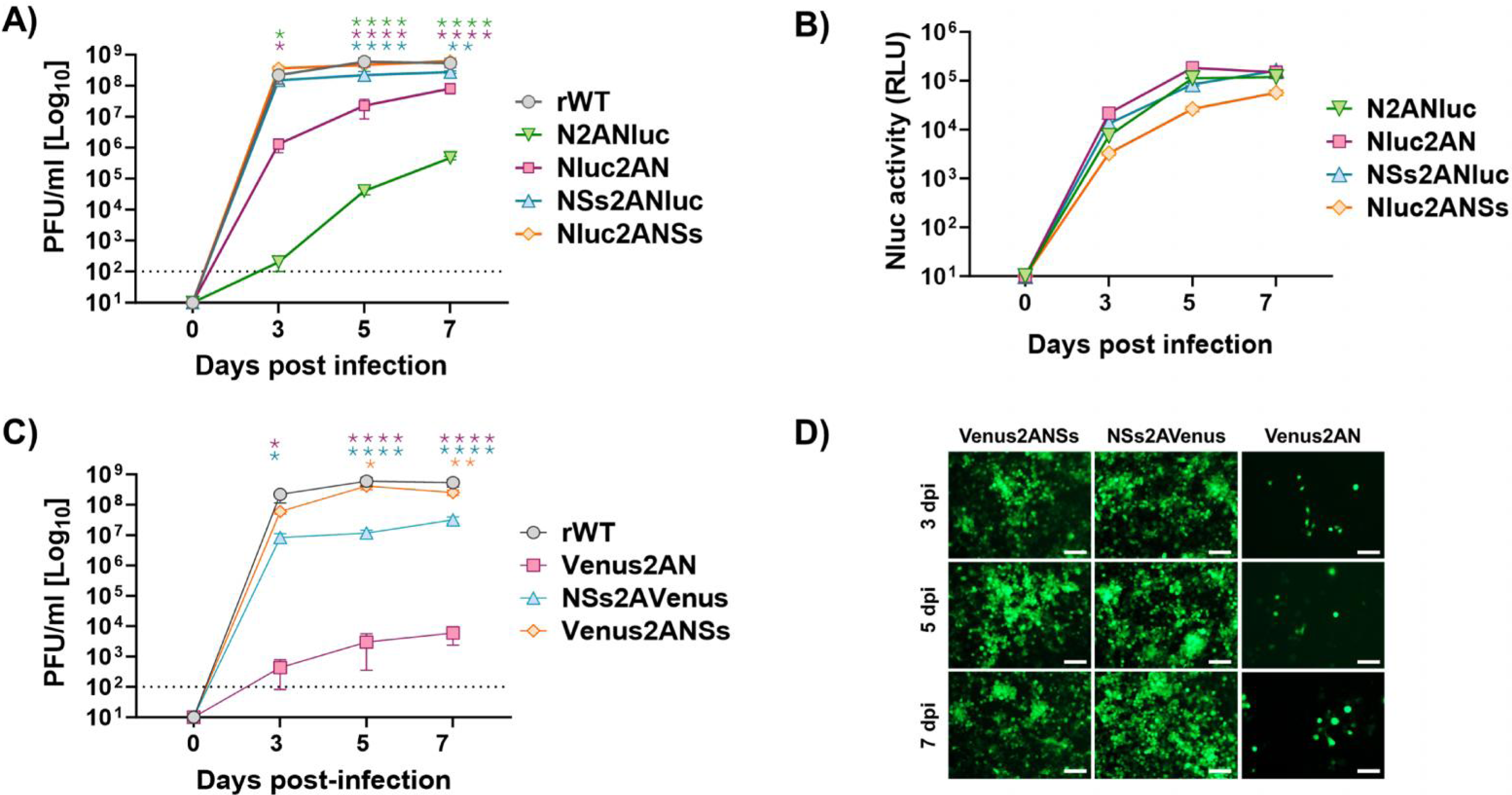
Multicycle growth kinetics in insect cells. C6/36 mosquito cells (T25 flasks, triplicates) were infected (MOI of 0.01) with rRVFV WT (rWT) or with recombinant viruses expressing Nluc (**A and B**) or Venus (**C and D**). At the indicated days p.i. (3, 5, and 7), viral titers in culture supernatants were evaluated by plaque assay (PFU/ml), and the same rWT representation was used in A and C. Dunnett’s method was used to statistically compare viruses expressing Nluc or Venus versus rWT. * P < 0.05; ** P < 0.0021; *** P < 0.0002; **** P < 0.0001 (**A and C**). The data represent the means ± SDs of triplicate samples. At the same time, p.i. Nluc activity (RLU: Relative Light Units) was measured in (**B**), and Venus expression was evaluated using a fluorescence microscope (**D**). Representative images are shown. Scale bars, 100 μm. Dotted line indicates the limit of detection (100 FFU/ml) of the assay.

### Pathogenicity and replication of Nluc-expressing rRVFV in mice

Next, we evaluated the pathogenicity of Nluc-expressing rRVFV in BALB/C mouse model of infection (**Figure 10**). To that end, groups of mice (n = 5) were inoculated intraperitoneally with 10^3^, 10^4^ or 10^5^ PFU of each rRVFV and body weight loss (**Figure 10A**) and survival (**Figure 10B**) were monitored for 18 days. Compared with rWT-infected mice, Nluc-expressing rRVFVs were attenuated. With a lower dose of 10^3^ PFU, all mice inoculated with rWT lost weight and all of them succumbed to viral infection. However, only rRVFV NSs2ANluc displayed virulence level comparable to the rWT, and although moderate body weight loss occurred between days 7 and 9, four of the 5 mice died. When animals were infected with 10^4^ PFU, only two of five mice inoculated with rWT died, and none of the mice infected with the Nluc-expressing rRVFVs showed signs of morbidity and all of them survived viral infection. Interestingly, with the higher dose of 10^5^ PFU, three animals inoculated with rWT and 2 inoculated with rRVFV Nluc2ANSs succumbed to the infection, although without significant body weight loss. The absence of a correlation between the dose and virulence was somehow expected, since this phenomenon has been shown before for RVFV (53, 54), including some of our studies (data not shown).

**Figure 10.**
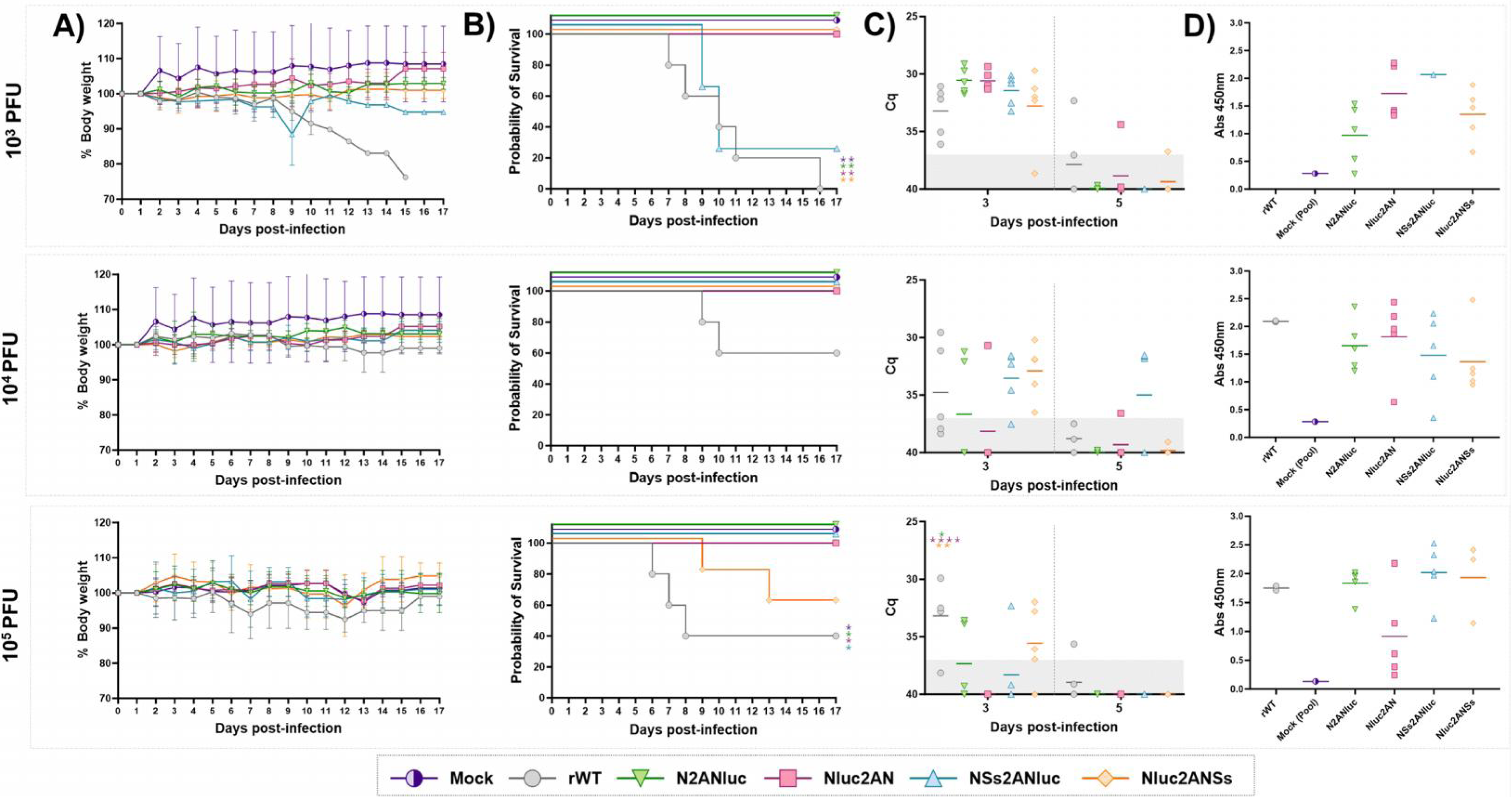
In vivo virulence of Nluc-expressing rRVFV. Groups of 8-to-12-week-old BALB/C female mice (N= 5/group) were infected with 10^3^, 10^4^ or 10^5^ PFU/mouse of the indicated viruses: rWT or the Nluc-expressing viruses. Weight loss (**A**) and survival (**B**) were evaluated daily for 18 days. The statistical differences between the curves of recombinant viruses expressing NLuc versus rWT were calculated using the log-rank test, with a P-value < 0.05. (**C**) Viremia determined by RT-qPCR in EDTA-treated blood samples collected at day 3 and 5 p.i. Cutoff: Cq ≥ 37 (dotted grey line). Points represent individual Cq value for each mouse, and lines of the corresponding colour represent the mean Cq value of each group. Dunnett’s method was used to statistically compare viruses expressing Nluc versus rWT. * P < 0.05; ** P < 0.0021; *** P < 0.0002; **** P < 0.0001. (**D**) Antibody responses in surviving mice were evaluated by ELISA (using purified N protein) on day 18 p.i., to confirm seroconversion. Mouse sera were diluted 1:100.

Given that RVFV is isolated from blood and causes viremia, it was speculated that the presence of Nluc could be detected in blood samples from infected mice, as occurs with Nluc-expressing bluetongue virus, which is also isolated from blood samples of infected mice (38). To prove this, Nluc activity was measured in plasma samples from infected animals and days 3 and 5 p.i. However, Nluc activity was not detected in any of the plasma samples (data not shown), even presence of the virus was detected at the same times post-infection by qRT-PCR (**Figure 10C**). Replication levels were higher at 3 days p.i in all cases and a clear correlation between mortality and leves of viral RNA in blood was not observed (**Figure 10C**). To study seroconversion in the animals, serum samples were collected at 18 d p.i, and the presence of antibodies against RVFV N was analyzed by ELISA (**Figure 10D**). The presence of anti-N antibodies is typically used as a sign of infection and replication and as a valid serological diagnostic method (20, 55–58). We observed seroconversion in all the mice that survived infection (**Figure 10D**), suggesting that Nluc-expressing rRVFVs were able to replicate, although they are mostly attenuated in vivo. Altogether, these data indicate that the S segment-based reporter gene expression strategy contributes to in vivo attenuation depending on the insertion position, at least in the mouse model of infection used.

## DISCUSSION

The establishment of reverse genetics systems for RVFV and other bunyaviruses has allowed researchers to investigate virus replication, mechanisms of viral pathogenesis, and the generation of live-attenuated vaccines (7, 17, 19, 45, 56, 59). Reporter-expressing, replication-competent rRVFV represents a powerful tool for basic and/or translational studies (9, 19, 22, 29). Previously, reporter-expressing rRVFVs were generated using a four segment strategy or by substituting the viral NSs sequence with the reporter gene (22, 29). In this work, we have assayed the insertion of a reporter gene at four different positions in the viral segment S, avoiding duplication of viral segments or removal of non-structural viral genes; and, therefore, representing a more biological approach than previous strategies. In this new approach, foreign sequences were expressed from the same viral N or NSs ORFs using the PTV-1 2A cleavage site, an approach that has been successfully implemented for the development of other replication-competent, reporter-expressing viruses with high success (38, 43, 44). Similar to these previous studies, we have been able to successfully rescue using this novel 2A approach RVFV expressing fluorescent and luciferase reporter genes. Based on the different properties of fluorescent and luciferases proteins (31–34, 36) it is important to carefully considered the advantages and disadvantages of these reporter genes before performing experiments. For example, *in vitro*, luciferases may be more appropriate for quantitative determinations (33, 35, 60). Conversely, fluorescent proteins are a better choice to observe localization in live cells (31, 32). Thus, in this manuscript we used the same innovative approach to generate rRVFVs expressing luciferase (Nluc) or fluorescent (Venus) reporter proteins. However, other reporter luciferase or fluorescent proteins can be inserted into the genome of RVFV using the same experimental strategy to rescue similar viruses. Moreover, based on the results obtained in this study, we have been able to have a better understanding on the plasticity of the RVFV genomic S segment to accommodate extra foreign sequences and how that influences expression levels of foreign proteins or viral attenuation of recombinant viruses expressing these reporter genes from the locus of the viral N or NSs ORFs.

Subsequently, the expression of the Nluc and Venus was confirmed, and the virological phenotypes of Nluc- and Venus-expressing viruses were characterized. As expected, the infection of mammalian Vero E6 or mosquito C6/36 cells with reporter-expressing rRVFV was visualized in real time without the need for secondary methods. Moreover, the expression of Venus and Nluc revealed similar kinetics that correlated with the levels of viral replication, further demonstrating the feasibility of using Nluc or Venus as valid surrogates of viral replication and infectivity without the need of secondary approaches to identify the presence of the virus in infected cells or cell culture supernatants, respectively. Compared to rWT, replication of the reporter-expressing rRVFVs was affected, although, in general, all viruses growth with high viral titers. Importantly, all the rRVFVs were stable during multiple passages in cell culture, which is desirable for studies using reporter-expressing viruses (36, 38, 41). However, we identified differences regarding the strategy used to insert the reporter genes (**Table 1**). Greater expression was observed when reporter genes were expressed from the locus of the viral N protein. On the other hand, rRVFV encoding Nluc or Venus from the locus of the NSs protein replicated to a greater extent but had reduced levels of reporter gene expression. Notably, if reporter genes were inserted after the UTR sequence, we observed higher viral titers than those of recombinant viruses expressing the reporter gene before the UTR sequence. However, the mechanisms responsible for these differences remain to be investigated and will be the subject of our follow up studies aimed to assess foreign gene expression from the RVFV S segment based on their location in the viral genome.

**Table 1.** Summary of strategies to generate reporter-expressing rRVFV.

| rRVFV | Properties |  |  |  |  |  |  |
| --- | --- | --- | --- | --- | --- | --- | --- |
|  | Growth (Vero E6) | Plaque phenotype | Growth (C6/36) | Expression Nluc/Venus | In vitro stability | MN/antiviral assay | Virulence |
| rWT | +++ | +++ | +++ | - | ND | +++ | +++ |
| Nluc2ANSs | +++ | +++ | +++ | ++ | ++ | +++ | + |
| NSs2ANluc | ++ | +++ | +++ | ++ | +++ | +++ | + |
| Nluc2AN | ++ | ++ | ++ | +++ | ++ | +++ | - |
| N2ANluc | + | + | + | +++ | +++ | ND | - |
| Venus2ANSs | +++ | +++ | +++ | +++ | ++ | ND | ND |
| NSs2AVenus | ++ | ++ | ++ | +++ | +++ | ND | ND |
| Venus2AN | ++ | + | + | +++ | +++ | ND | ND |

ND: Not determined.

Notably, we developed and validated a Nluc-based microneutralization assay that allow us to rapidly identify antiviral compounds and NAbs against RVFV. Our novel reporter-based mincroneutralization assay represent an excellent option to interrogate large libraries of compounds or antibodies to identify those with antiviral and neutralizing activity against RVFV, including their feasibility to implement in high-throughput (HTS) approaches to rapidly and sensitivity screen for novel antivirals or, in the case of NAbs, the efficacy of novel vaccine candidates. In addition, the virulence of Nluc-expressing rRVFVs was assessed in BALB/C mice. In this mouse model of infection, reporter-expressing viruses were attenuated as compared to rWT, and mortality was only observed in mice inoculated with Nluc2ANSs or NSs2ANluc viruses. Importantly, all reporter rRVFVs were able to replicate in this mouse model. However, the reporter rRVFVs could display a more virulent phenotype in other mouse strains that are more susceptible to RVFV infection (61). It has been shown that BALB/C mice (used in our study) are slightly more resistant than other mouse strains, where different disease phenotypes (hepatic, neurological and mixed) can be clearly distinguished (53, 54, 62). In addition, differences depending on the sex and age of the animals have been reported, with males being more susceptible than females (63, 64). However, in our study, we have used only female BALB/C mice, and the phenotype of our rRVFV in other mouse models or in male vs female animals remains unknown. Interestingly, all mice produced anti-N antibodies, suggesting that the viruses were able to infect and replicate (20, 55–58), although causing limited or no pathogenicity in terms of body weight loss or mortality. Altogether, our results indicate that additional studies to determine the best mouse strain, sex and viral dose should be taken into account when using the generated reporter-expressing rRVFV.

NSs is localized both in the cytoplasm and nucleus of infected cells and it is considered the major virulence factor of RVFV. NSs form nuclear filamentous structures through homo-oligomerization (65) and inhibit host innate immune responses through different strategies, including inhibition of the type I interferon (IFN) system (66), and induction of host cellular transcription shut-off (67). It has been suggested that NSs filament formation is dispensable for efficient replication in vitro and appears to be important but insufficient for virulence in vivo (68). In two of our strategies used to generate reporter-expressing rRVFVs, we introduced the foreign sequence adjacent to the NSs sequence. This could affect some of the antiviral functions of NSs, including the formation of filamentous structures, and thus explain the attenuated phenotype in vivo. However, filament like structures resembling those of WT NSs were clearly observed in the virus expressing NSs2ANluc and to a lesser extent in the Nluc2ANSs virus. Interestingly, the detection of such filaments was accomplished using an anti-Nluc antibody, indicating that unprocessed NSs-Nluc or Nluc-NSs fused proteins were transported to the nucleus of the infected cells. How this fact correlates with the observed virulence levels in vivo of both rRVFVs remains to be elucidated. In addition, it could be interesting, to compare the viral fitness and attenuation of the described rRVFV encoding the reporter gene in the NSs side with a rRVFV lacking the viral NSs gene.

The formation of the RVFV N-RNA complex protects the vRNA and replication intermediates from nucleases and is key for viral replication-transcription, and efficient packaging of the viral genome (69, 70). It has been observed in crystallographic structures that the N protein adopts two major conformations, open and closed (69, 71–73). It cannot be discarded that by inserting the reporter gene next to the viral N sequence, the proper function of the N protein during viral infection could be affected, due to the intrinsic proteolytic mechanism of the PTV-2A peptide in which the cleavage site lies between the Gly-Pro bond at the end of the peptide, therefore 21 extra amino acids are added to the end of the viral nucleoprotein. This could explain the observed attenuation in vivo and reduced viral fitness in vitro compared with those of the rWT or with rRVFVs expressing the reporter gene on the NSs side.

In addition, although the UTRs and IGR of the viral segment S were not directly modified in our strategy, the insertion of foreign sequences could introduce some changes in the RNA structure or unknown cis-regulatory sequences of the viral genomic and/or antigenomic sequences. In fact, it was observed, that reporter-expressing rRVFVs containing the reporter gene next to the 5’ or 3’ UTRs displayed better fitness than viruses in which the heterologous genes were inserted next to the IGR (17, 74–76). Notably, our strategy could also be useful for developing and testing novel live RVFV vaccines or vaccine vectors with different degrees of attenuation.

## MATERIALS AND METHODS

### Cell lines and viruses

Vero E6 cells (ATCC/CRL/1586) were routinely maintained in Dulbecco’s modified Eagle’s medium (DMEM) supplemented with 5% fetal bovine serum (FBS), 1% 100x nonessential amino-acids, 100 U/mL penicillin, 100 μg/mL streptomycin, and 2 mM L glutamine. The cells were incubated at 37°C in the presence of 5% CO_2_. BSR-T7 cells (38, 77) stably expressing the bacteriophage T7 RNA polymerase were maintained in the same manner as above except for the addition of 1 mg/mL of Geneticin every 2-3 passages. C6/36 *Aedes albopictus* cells (ATCC CRL–1660) were grown in Eagle’s Minimum Essential Medium (MEM) supplemented with 10% fetal calf serum (FCS), L-glutamine (2 mM), gentamicin (50 μg/ml), and MEM vitamin Solution (Sigma) at 28°C.

The sequence of the WT South African RVFV strain 56/74 (parental virus), which was used for the synthesis of the reverse genetic system, was previously described (78). GenBank: OM744400.1 (segment L), OM744401.1 (segment M) and OM744402.1 (segment S). Virus stocks were propagated in Vero E6 cells and titrated by plaque assay as previously described (20, 55, 56). All the experiments were conducted in the biological security facility of the Animal Health Research Center (CISA, INIA-CSIC).

### Plasmids

All pBlueScript II SK (+) reverse genetics (RG) plasmids were synthetized *de novo* by Biomatik and designed to contain the viral cDNAs flanked upstream by a T7 RNA polymerase promoter sequence and downstream by the hepatitis delta virus ribozyme (HDVR) and T7 RNA polymerase terminator sequences (38). To generate replication-competent reporter-expressing RVFVs and study viral genome plasticity, we engineered a set of recombinant modified segments S using four different strategies for the cloning of reporter genes. Importantly, the 5′- and 3′-UTRs and the IGR were not modified. A common heterologous sequence was designed containing a linker (GSG), the porcine teschovirus-1 (PTV-1) 2A autoproteolytic cleavage site (ATNFSLLKQAGDVEENPGP), and a multicloning site (AgeI/BglII/NheI) for the insertion of foreign sequences (36, 38, 42). Then, the 4 master plasmids containing the heterologous sequence before or after the 5′- and 3′-UTRs or the IGR were used for the cloning of the Venus or Nluc reporter genes. Reporter genes were amplified by PCR using specific primers containing AgeI or NheI sites. Then, the PCR products and the 4 modified segment S plasmids were digested, and the reporter genes were cloned using standard molecular biology techniques. Plasmid constructs were confirmed by sequencing (Macrogen). Master plasmids containing the modified segment S

-pBS-2A-N: The reporter gene without a stop codon was cloned between the 3’UTR of the genomic vRNA and the N ORF. This plasmid was used to generate Venus2AN and Nluc2AN viruses.

-pBS-N-2A: The reporter gene with a stop codon was cloned between the N ORF and the IGR. This plasmid was used to generate N2AVenus and N2ANluc viruses.

-pBS-NSs-2A: The reporter gene with a stop codon was cloned between the NSs ORF and the IGR. This plasmid was used to generate NSs2AVenus and NSs2ANluc viruses.

-pBS-2A-NSs: The reporter gene without a stop codon was cloned between the 5’UTR of the genomic vRNA and the NSs ORF. This plasmid was used to generate Venus2ANSs and Nluc2ANSs viruses.

### Rescue of recombinant viruses

To rescue rRVFV, monolayers of BSR-T7 cells (T25 flasks, 80% of confluence, duplicates) were cotransfected with 1 µg of each of the reverse genetics plasmids encoding the RVFV cDNAs using Lipofectamine 3000 (Invitrogen) in Opti-MEM I reduced serum medium (Invitrogen) according to the manufacturer’s instructions. Twenty-four hours after transfection, the transfection mixture was replaced with 5 mL of growth medium. After 72-96 h, the cell culture supernatants were collected, clarified, and used to infect fresh monolayers of Vero E6 cells. Once the CPE was observed (48-96 h), the medium was harvested and clarified. Clonal rRVFV were selected by plaque assays, and virus stocks were propagated in Vero E6 cells, where once the CPE was observed, the medium was harvested, clarified and stored at −80°C.

### Virus growth kinetics

To evaluate virus multicycle growth kinetics, confluent Vero E6 cells (12-well plate format, 5 x 10^5^ cells/well, triplicates) or C6/36 cells (T25 flasks, 4 x 10^6^ cells/well, triplicates) were infected at an MOI of 0.01. After 1 h of virus adsorption at 37°C (Vero E6) or 28°C (C6/36), the inoculum was discarded, the cells were washed, and 1.5 mL (Vero E6) or 5 mL (C6/36) of growth medium was added. At the indicated times post infection (24, 48, and 72 h for Vero E6; 3, 5, and 7 days for C6/36), 150 µL of the supernatant was removed and stored at −80°C. Virus titers were determined in Vero E6 cells by plaque assay (plaque-forming units, PFU/mL), as previously described (20, 55, 56, 78). The mean value and standard deviation (SD) were calculated using GraphPad Prism version 8.0.1 (GraphPad Software Inc.). In addition, cells infected with Venus-expressing rRVFV were imaged using a Zeiss Axio fluorescence microscope (Zeiss), and the presence of Nluc in the culture supernatants was quantified using Nano-Glo luciferase substrate (Promega) following the manufacturer’s specifications.

### Plaque assay and immunostaining

Confluent monolayers of Vero E6 cells (6-well plate format, 10^6^ cells/well) were infected with the indicated viruses for 1 h at room temperature, overlaid with agar, and incubated at 37°C. At 3 days p.i., the cells were fixed with 10% paraformaldehyde (PFA), and the overlays were removed. The cells were then permeabilized (0.01% Triton X-100 in PBS) for 15 min at room temperature, and prepared for immunostaining as previously described (36). Primary monoclonal or polyclonal antibodies (mAbs and pAbs, respectively) antibodies against Nluc (mouse mAb; R&D Systems) for Nluc-expressing viruses, GFP (goat pAb, Rockland) for Venus-expressing viruses, and anti-N (mAb, 2B1) (59) were used. The Vectastain ABC kit and DAB HRP substrate kit; (Vector) were used for the immunostaining following the manufacturers’ specifications. The plates were stained with crystal violet after the immunostaining.

### Protein gel electrophoresis and Western blot analysis

Cell extracts from either mock- or virus-infected (MOI, 0.1) Vero E6 cells were lysed at 24 or 48 h p.i. in radioimmunoprecipitation assay (RIPA) buffer, and proteins were separated by denaturing electrophoresis. The membranes were blocked for 1 h with 5% dried skim milk in TBS containing 0.1% Tween 20 (T-TBS) and incubated overnight at 4°C with specific primary mAbs and pAbs: Nluc (mouse mAb; R&D Systems) for Nluc-expressing viruses, GFP (goat pAb, Rockland) for Venus-expressing viruses, anti-PKR (rabbit pAb, Invitrogen), NSs (pool of mAb; 5C3A1B2, 2A10B1B and 5F12C1B3) kindly provided by M. Groschup (45), and anti-RVFV (rabbit pAb generated in the laboratory, using purified virus and detecting viral Gn/Gc and N proteins) (20). A mAb against actin (Sigma) was used as an internal loading control. Bound primary antibodies were detected with horseradish peroxidase (HRP)-conjugated secondary antibodies against the different species (mouse, rabbit or goat). Proteins were detected by chemiluminescence (Thermo Fisher Scientific) following the manufacturer’s recommendations and photographed using a Bio-Rad digital ImageStation.

### Indirect immunofluorescence assays

Vero E6 cells were mock infected or infected (MOI, 0.1) with the indicated rRVFV. At 24 h p.i., the cells were fixed with 4% paraformaldehyde (PFA) and permeabilized with 0.5% Triton X-100 in PBS for 15 min at room temperature and blocked with PBS/5% FBS. After blocking, specific primary mAbs or pAbs for Nluc (mouse mAb; R&D Systems) for Nluc-expressing viruses, GFP (goat pAb, Rockland) for Venus-expressing viruses, and anti-RVFV (rabbit pAb generated in the laboratory using purified virus that detects the viral Gn/Gc and N proteins) (20) were diluted in blocking solution and used to separately detect protein expression using Alexa Fluor 594-conjugated secondary antibodies (Invitrogen). Images were taken with a Zeiss Axio fluorescence microscope (Zeiss) or with a Zeiss LSM880 confocal laser microscope (Zeiss).

### Flow cytometry assays

Vero E6 cells were mock infected or infected (MOI, 0.01) with the indicated rRVFV WT or Venus-expressing viruses. At 24 h p.i., the cells were harvested from the culture, centrifuged for 2 min at 2,000 x g and incubated for 30 min at 4°C with a mAb against the N (2B1) conjugated to Alexa 700 (Invitrogen). The cells were washed and fixed with fixation buffer (BioLegend) for 20 minutes at room temperature in the dark. The cells were analyzed by flow cytometry by gating on green (Venus) and red fluorescence (mAb anti-N 2B1 Alexa 647 conjugated) (59) using a CyFlow Cube 8 cytometer (Sysmex). The data were analyzed using FlowJo v10 software.

### Stability of reporter-expressing viruses in cultured cells

After generation of virus working stocks (P2), confluent monolayers of Vero E6 cells (24-well plate format, duplicates) were infected (MOI, 0.01) with the reporter expressing viruses and incubated until a 70% CPE was observed. Tissue culture supernatants were then harvested and 1/10 of the collected medium was used for the infection of fresh Vero E6 cells for a total of 3 passages. During each viral passage, cultured supernatants were collected and plaque assays were carried out as described above. Venus-expressing plaques were visualized directly using a Zeiss Axio fluorescence microscope (Zeiss). To determine the stability of Nluc-expressing viruses, plaques were used to infect fresh Vero E6 cells (96-well plate format, 2.5 x 10^4^ cells/well), and Nluc activity in the tissue culture supernatants was determined at 48 h.p.i., as indicated above. Approximately 30 plaques per passage were evaluated for each virus.

### Reporter-based microneutralization assays

Nluc microneutralization assays were performed as previously described (36–38). Pre- and post-challenge Sheep sera from a previous experiment carried out in the laboratory were serially diluted 2-fold (starting dilution, 1/20). Then, 200 PFU of virus was added to the antibody dilutions, and the mixture was incubated for 1 h at 37°C. Vero E6 cells (96-well plate format, 5 x 10^4^ cells/well, triplicates) were then infected with the serum-virus mixture for 1 h at 37°C. Then, the serum-virus mixture was removed and 150 µl of fresh growth medium was added and incubated for 48 h at 37°C. The Nluc activity in the culture supernatants was quantified using Nano-Glo luciferase substrate (Promega) and a microplate reader as described previously. The luciferase values of virus-infected cells in the absence of serum were used to calculate 100% viral infection. Cells in the absence of viral infection were used to calculate the luminescence background. Triplicate wells were used to calculate the average and SD of neutralization. The 50% neutralization titter (NT_50_) was determined by use of a sigmoidal dose-response curve (GraphPad Prism, v8.0.1 software).

### Antiviral assays

Confluent monolayers of Vero E6 cells (96-well plate format, 5 x 10^4^ cells/well, triplicates) were infected with 200 PFU of the Nluc-reporter rRVFVs. After 1 h of infection, the fresh medium was supplemented with 2-fold serial dilutions (starting concentration, 1000 μM) of ribavirin (TCI), and the cells were incubated for 48 h at 37°C. Then, Nluc expression was quantified as indicated above. Triplicate wells were used to calculate the mean and SD of inhibition. The 50% inhibitory concentration (IC_50_) was determined by use of a sigmoidal dose-response curve (GraphPad Prism, v8.0.1 software).

### Mouse experiments

The animals were housed under pathogen-free conditions at the biosafety level 3 (BSL3) animal facility of the Animal Health Research Center (CISA-INIA/CSIC), Madrid (Spain). Animal experimental protocols were approved by the Ethical Review Committee at the CISA-INIA/CSIC and Comunidad de Madrid (Permit number: PROEX079.6/22), in strict accordance with EU guidelines 2010/63/UE about the protection of animals used for experimentation and the Spanish Animal Welfare Act 32/2007. For pathogenicity studies, cohorts of 8-to-12-week-old BALB/C female mice (N = 5/group) were intraperitoneally inoculated with the indicated viruses at the indicated dose. Then, the animals were monitored daily for 18 days for clinical symptoms (such as malaise, respiratory distress, and lack of movement), body weight loss relative to the initial weight, and mortality. At 72 or 96 h after infection, blood samples were taken by submandibular puncture and tested for viral RNA by RT-qPCR (59, 79) to monitor viremia. To assess the presence of Nluc in blood samples, plasma from 50 µL of the same whole blood sample was obtained, and the Nluc activity was measured using a Nano-Glo Luciferase Assay Kit (Promega) and a microplate reader (38). Serum samples to be used in antibody assays were collected at 18 d.p.i. (at the end of the experiment) by submandibular puncture. Due to space limitations in the BSL3 facility, experiments using 10^5^ and 10^3^/10^4^ were conducted in different days/weeks. However, the same viral stocks and mouse provider were used. All the inoculums were prepared via serial dilution to limit variability in the dose, and the virus was back tittered after mouse inoculation to discard mistakes.

### ELISA

Antibodies against N were detected by an in-house ELISA (55). Briefly, for the detection of antibodies against N, sera collected at 18 d.p.i were tested (Dilution 1/100) in ELISA plates adsorbed with 50 ng/well of purified recombinant N protein produced from *Escherichia coli* and diluted in carbonate buffer (pH 9.6). The wells were blocked with 5% BSA in PBS and 0.05% Tween 20 (PBS-T). Then, the bound antibodies were detected with conjugated goat anti-mouse-IgG-HRP, and the bound conjugates were detected using TMB for 10 min, followed by the addition of one volume of stopping solution (3N H_2_SO_4_). Optical densities were measured at 450 nm (OD450).

### Statistical analysis

Microsoft Excel (Microsoft Corporation) and GraphPad Prism version 10.1.2 (GraphPad Software, San Diego, CA, USA) software were used to analyze the data. Microsoft Excel was used to perform some of the calculations and to visualize the raw data. GraphPad Prism software was used to perform statistical analysis of all the data. Growth kinetics were analyzed using two-way ANOVA the Dunnett’s multiple comparisons test. A one-way ANOVA with Tukey’s method was used to evaluate viral titters comparing between groups. To evaluate the Nluc activity between the different groups, two-way ANOVA by Tukey’s method was used. Statistical differences between survival curves were analyzed using log-rank test. Viremia levels were analyzed using the Dunnett’s method. A P value lower than 0.05 was considered significant in all cases.

## DATA AVAILABILITY STATEMENT

All data supporting the findings of this study are available within the paper. No datasets or new viral sequences were generated or analysed during the current study.

## ACKNOWLEDGEMENTS

This study was partially funded by the Ramon y Cajal Incorporation grant (RYC-2017) from the Spanish Ministry of Science, Innovation, and Universities and the European Uniońs Horizon Europe Research and Innovation Programme (grant agreement 101046133), and by grant PID2021-122567OB-I00 funded by MCIN/ AEI /10.13039/501100011033/ and by ERDF “A way of making Europe”.

